# A 1-bp deletion in bovine *QRICH2* causes low sperm count and immotile sperm with multiple morphological abnormalities

**DOI:** 10.1101/2021.11.19.469233

**Authors:** Maya Hiltpold, Fredi Janett, Xena Marie Mapel, Naveen Kumar Kadri, Zih-Hua Fang, Hermann Schwarzenbacher, Franz R Seefried, Mirjam Spengeler, Ulrich Witschi, Hubert Pausch

## Abstract

**Background:** Semen quality and male fertility are monitored in artificial insemination bulls to ensure high insemination success rates. Only ejaculates that fulfill minimum quality requirements are processed and eventually used for artificial inseminations. We examined 70,990 ejaculates from 1343 Brown Swiss bulls to identify bulls from which all ejaculates were rejected due to low semen quality. This procedure identified a bull that produced twelve ejaculates with an aberrantly low number of sperm (0.2±0.2 × 10^9^ sperm per ml) which were mostly immotile due to multiple morphological abnormalities.

**Results:** The genome of the bull was sequenced at 12-fold coverage to investigate a suspected genetic cause. Comparing the sequence variant genotypes of the bull with those from 397 fertile bulls revealed a 1-bp deletion in the coding sequence of *QRICH2* encoding glutamine rich 2 as a compelling candidate causal variant. The 1-bp deletion causes a frameshift in translation and induces a premature termination codon (ENSBTAP00000018337.1:p.Cys1644AlafsTer52). The analysis of testis transcriptomes from 76 bulls showed that the transcript with the premature termination codon is subjected to nonsense-mediated mRNA decay. The 1-bp deletion resides on a 675 kb haplotype spanning 181 SNPs from the Illumina BovineHD Bead chip. The haplotype segregates at a frequency of 5% in the Brown Swiss cattle population. This analysis also identified another bull that carried the 1-bp deletion in the homozygous state. Semen analyses from the second bull confirmed low sperm concentration and immotile sperm with multiple morphological abnormalities primarily affecting the sperm flagellum and, to a lesser extent, the sperm head.

**Conclusions:** A recessive loss-of-function allele of bovine *QRICH2* likely causes low sperm concentration and immotile sperm with multiple morphological abnormalities. Routine sperm analyses unambiguously identify homozygous bulls. A direct gene test can be implemented to monitor the frequency of the undesired allele in cattle populations.

## Background

Semen quality and male fertility are complex traits with low to moderate heritabilities [1]. These traits are routinely monitored in thousands of bulls as a service from the semen collection centres to the cattle breeding industry to ensure high success rates in artificial insemination. Large cohorts with repeated semen quality analyses facilitate separating genetic from environmental factors as well as differentiating between female and male factors contributing to variation in reproduction [2]. Such phenotypes are ideally suited to characterize the genetic architecture of male fertility [3].

Genome-wide association testing revealed quantitative trait loci (QTL) for semen quality and male fertility in various breeds of cattle, many of them expressing non-additive effects [4],[5],[6]. The routine evaluation and recording of tens-of thousands of ejaculates and millions of artificial inseminations also occasionally detect bulls with aberrant semen quality or strikingly low insemination success rates. Although diseases and environmental exposures can compromise male fertility [7], drastically reduced semen quality in individual bulls often results from monogenic disorders [8],[9],[10],[11]. Genome-wide case-control association testing facilitates pinpointing variants for such conditions. In this approach, genotypes at genome-wide markers are compared between affected bulls and an unaffected control group. Variants at which the genotypes differ between both groups are candidate causal variants that are subsequently subjected to further functional investigations.

Impaired semen quality and low fertility in healthy bulls can result from deleterious alleles in genes that show testis-biased or testis-specific expression [8],[10],[9]. Such alleles are spread disproportionately by females because they are typically not impacted by defective genes that are primarily expressed in the male reproductive system. Semen quality and male fertility are only evaluated in breeding bulls. Thus, recessive variants that compromise male reproduction can remain undetected in cattle populations for a long period of time or reach high allele frequency without being recognized [8].

Here we report a frameshift-inducing 1-bp deletion in bovine *QRICH2* that likely causes male infertility in the homozygous state due to immotile sperm with multiple morphological abnormalities. Using historic and current semen quality data from two Brown Swiss bulls, we show that this allele remained undetected although phenotypic consequences became manifest almost two decades ago. Our findings offer the opportunity to implement direct gene testing to monitor the frequency of the undesired allele in cattle populations.

## Material and Methods

### Animals and phenotypes

We assessed records for 70,990 ejaculates that were collected from 1343 Brown Swiss bulls between January 2000 and March 2018. All ejaculates were examined by laboratory technicians employed by Swissgenetics immediately after semen collection as part of routine quality assurance during semen processing. Parameters recorded were semen volume (in ml), sperm concentration (million sperm per ml) quantified using photometric analysis, and the percentage of sperm with forward motility. The proportion of sperm with head and tail anomalies was quantified for each ejaculate with scores ranging from 0 to 3 (0: no or very few anomalies, 1: less than 10% sperm with anomalies, 2: between 10 and 30% sperm with anomalies, 3: more than 30% sperm with anomalies). We subsequently considered 35,785 ejaculates that were collected from 1258 bulls at the age between 330 and 550 days to identify bulls from which all ejaculates were discarded due to insufficient semen quality. Ejaculates that contained less than 300 million sperm per ml, less than 70% motile sperm, more than 10% sperm with head and tail abnormalities, or whose volume was less than 1 ml were deemed to be unsuitable for artificial inseminations.

### Whole-genome sequencing and sequence variant genotyping

We extracted DNA from hair roots of a Brown Swiss bull (sequence read archive accession: SAMEA6272098) that produced ejaculates with low sperm count and immotile sperm with multiple morphological abnormalities. For whole-genome sequencing, an Illumina TruSeq DNA PCR-free paired-end library with 400 bp insert size was sequenced at an Illumina NovaSeq6000 instrument. We used the fastp software [12] to remove adapter sequences, poly-G tails and reads that had phred-scaled quality less than 15 for more than 15% of the bases. Following quality control, we aligned 153,618,844 read pairs to the ARS-UCD1.2 assembly of the bovine genome using the mem-algorithm of the BWA software [13] with option -M to mark shorter split hits as secondary alignments and default settings for all other parameters. We marked duplicates using the Picard tools software suite (https://github.com/broadinstitute/picard). Alignments were sorted by coordinates using Sambamba [14]. Sequencing depth was calculated using the mosdepth software (version 0.2.2, [15]) considering only reads with mapping quality >10.

Sequence variants (SNPs and indels) were genotyped for SAMEA6272098 together with 521 cattle from various breeds (14 Hereford, 1 Nelore, 33 Grauvieh, 50 Fleckvieh, 2 Nordic Red Cattle, 47 Holstein, 243 Brown Swiss, 128 Original Braunvieh, 3 Wagyu) using a multi-sample variant discovery and genotyping approach implemented with the HaplotypeCaller, GenomicsDBImport and GenotypeGVCFs modules from the Genome Analysis Toolkit (GATK, [16]). Subsequently, we applied best practice guidelines of the GATK for variant filtration and imputed missing genotypes using Beagle [17]. The reference-guided variant genotyping workflow applied here is described in detail in Crysnanto et al. [18]. Functional consequences of 41,659,308 autosomal variants were predicted according to the Ensembl (version 104) and Refseq (version 106) annotations of the bovine genome using the Variant Effect Predictor software [19] from Ensembl along with the SpliceRegion.pm plugin (https://github.com/Ensembl/VEP_plugins/blob/release/101/SpliceRegion.pm).

### Identification of candidate causal variants

We considered 397 sequenced fertile bulls as controls to identify sequence variant genotypes associated with the sperm disorder of SAMEA6272098 (**Additional file 1 File S1**).

Assuming recessive inheritance of a deleterious allele, we screened the sequence variant genotypes of SAMEA6272098 for homozygous sites that were never found in homozygous state in fertile bulls. These variants were subsequently filtered to retain those with a predicted deleterious impact on a protein.

### Identification of haplotype carriers in the population

Sequence variant and partially imputed Illumina BovineHD genotypes, respectively, of 285 and 33,045 cattle were used to assign the 1-bp deletion onto a haplotype. Briefly, 33,045 cattle were genotyped using different Illumina SNP microarrays [4]. Following quality control, the genotypes were imputed to a density of 683,903 SNPs with Beagle [20] using a reference panel of 1166 cattle that had BovineHD genotypes. We retained 25,243 Brown Swiss and 5,228 Original Braunvieh animals from the Swiss populations.

We inspected the phased (and partially imputed) BovineHD genotypes of 285 animals which also had genotypes for the 1-bp deletion called from whole-genome sequencing data.Specifically, we searched for a haplotype extending BTA19:55436705 on either side that was shared in the heterozygous state in all sequenced 1-bp deletion carriers. The genotypes and haplotypes of the 285 animals are given in **Additional file 1 File S1**. We used the alleles of the longest shared haplotype to identify heterozygous and homozygous haplotype carriers among 30,471 genotyped Brown Swiss and Original Braunvieh cattle. Alleles and positions of markers encompassed by the haplotype (**Additional file 2 File S2**) were also provided to Brown Swiss breeding associations and genetic evaluation centres to screen their genomic prediction reference populations and selection candidates for homozygous haplotype carriers.

### Whole transcriptome sequencing and read alignment

We used DNA and RNA sequencing data from a previously established expression QTL (eQTL) cohort to investigate transcript abundance and functional consequences arising from the 1-bp deletion. The eQTL cohort consisted of 76 mature bulls from which testes were sampled at a commercial slaughterhouse after regular slaughter. Total RNA and DNA was extracted from testis tissue and prepared for sequencing as described earlier [21].

DNA samples were sequenced on an Illumina NovaSeq6000 sequencer using 150 bp paired-end libraries. Quality control (removal of adapter sequences and bases with low quality, and trimming of poly-G tails) of the raw sequencing data was carried out using the fastp software [12] with default parameters. Following quality control, between 70,493,763 and 307,416,205 read pairs per sample were aligned to the ARS-UCD1.2 version of the bovine reference genome [22] using the mem-algorithm of the BWA software (see above). Average coverage of the 76 DNA samples estimated using the mosdepth software (see above) ranged from 6.3 to 27.6 with a mean value of 12.6 ± 4.2. Sequence variant genotypes were called and filtered using the HaplotypeCaller, GenomicsDBImport, GenotypeGVCFs and VariantFiltration modules from the GATK as described above.

Total RNA sequencing libraries (2×150 bp) were prepared using the Illumina TruSeq Stranded Total RNA sequencing kit and sequenced on an Illumina NovaSeq6000 sequencer. Quality control (removal of adapter sequences and bases with low quality, and trimming of poly-A and poly-G tails) of the raw sequencing data was carried out using the fastp software [12]. Following quality control, between 191,160,837 and 386,773,085 filtered reads per sample (mean: 283,587,831 ± 43,284,185) were aligned to the ARS-UCD1.2 reference sequence and the Ensembl gene annotation (release 104) using STAR (version 2.7.9a) [23] with options --twopassMode Basic, --sjdbOverhang 100, --outFilterMismatchNmax 3, and -- outSAMmapqUnique 60.

### Bioinformatic analysis

The structure of bovine *QRICH2* (ENSBTAG00000030173, Gene-ID 530282) was analysed using data from Ensembl (version 104) and Refseq (version 106). Transcript abundance (in transcripts per million, TPM) was quantified using kallisto [24] and aggregated to the gene level using the R package tximport [25]. Exon abundance was quantified using QTLtools [26]. Exon-exon junctions were visualized using the ggsashimi R package [27]. Read coverage as well as reference and alternative allele support at the 1-bp deletion were inspected using the Integrative Genomics Viewer (IGV, [28]).

### Genotyping of the 1-bp deletion

Polymerase chain reaction (PCR) and Sanger sequencing were used to validate the 1-bp deletion. PCR products from genomic DNA were amplified using the following primers: 5′- CATCGAGAAGGTGCAGATCC-3′ (forward) and 5′-CTGCCC ACCGTTTGTAGC-3′ (reverse) with a standard 50 µL PCR reaction mix (100 ng genomic DNA and final concentrations of 1X PCR reaction buffer, 200 µM dNTPs mixture, 0.2 µM each primer and 0.05 units/µL JumpStart Taq Polymerase (Sigma)). PCR amplification program consisted of the initial denaturation step at 94°C for 30s followed by 30 cycles of 94°C for 10s, 55°C for 1 min, and 72°C for 30s with a final extension step at 72°C for 1 min. All PCR products were purified with GenElute PCR Clean-Up Kit (Sigma) and sent for Sanger sequencing (Microsynth, Switzerland). Sequencing results were analysed using CLC Genomics Workbench software (Qiagen).

### Semen analyses in a bull homozygous for the 1-bp deletion

Ejaculates from a bull homozygous for the 1-bp deletion were collected at 14, 16, 17 and 20 months of age using an artificial vagina. Sperm concentration, total sperm count, and sperm motility were determined with an IVOS II CASA system (Hamilton Thorne Inc., Beverly, U.S.A.) using Leja 2-chamber slides (Leja, Nieuw-Vennep, the Netherlands). Semen was fixed in buffered formol saline solution (Na2HPO4 4.93g, KH2PO4 2.54g, 38% formaldehyde 125ml, NaCl 5.41g, distilled water q.s. 1000ml) and smears were prepared according to Hancock [29] for morphological examination. At least 200 spermatozoa were subsequently evaluated by phase contrast microscopy using oil immersion (Olympus BX50, UplanF1 100×/1.30, Olympus, Wallisellen, Switzerland). Morphologically abnormal sperm were assigned to defect categories according to Blom [30].

Sperm viability was assessed using the eosin-nigrosin staining method [31]. At least 200 spermatozoa were evaluated in stained slides under oil immersion using a light microscope (Olympus BX50, UplanF1 100×/1.30, Olympus, Wallisellen).

The bull was slaughtered at 21 months at a regular slaughterhouse. Testis tissue was preserved in formalin, embedded in paraffin and subsequently stained with hematoxylin-eosin for histological evaluation.

### Transmission electron microscopy

Sperm were prepared for transmission electron microscopy (TEM) from fresh semen diluted in OptiXcell (IMV technologies, L’Aîgle, France) and stored at 5°C overnight. Semen samples were washed twice by dilution in phosphate buffered saline (PBS) and subsequent centrifugation at 300g for 10 minutes. After removing the supernatant, the pellet was fixed with equal volume of 6% glutaraldehyde in PBS, resuspended gently, and centrifuged at 6000g. After removing the supernatant, the sperm were fixed for a second time with 3% glutaraldehyde in PBS, and finally pelleted at 6000g. Pellets were washed three times in PBS, post-fixed in 1% osmium tetroxide, washed in double distilled water, stained in 1% uranyl acetate, dehydrated in graded series of ethanol (25, 50, 75, 90, and 100%), and embedded in epoxy resin through increasing concentrations (25, 50, 75, and 100%) using PELCO Biowave+ tissue processor (Ted Pella, USA), and then cured at 60°C for three days. Embedded blocks were sectioned using Leica FC6 microtome and a DIATOME diamond knife with 45° angle into 60nm sections and mounted on Quantifoil copper grids with formvar carbon films. Sections were post-stained with 2% uranyl acetate followed by lead citrate. Grids were imaged using FEI Morgagni 268 electron microscope operated at 100kV at 8.9 to 56k magnification.

### Scanning electron microscopy

Sperm were prepared for scanning electron microscopy (SEM) within 30 minutes after semen collection. A thin layer of native semen was spread on carbon coated coverslips with the side of a pipet tip. The coverslips were immersed in 3% glutaraldehyde in PBS solution and kept on ice. After fixation overnight at 4°C, fresh fixative was added and the sample was put in a PELCO BioWave, Pro+ microwave system (Ted Pella, USA). Following the microwave-assisted fixation and dehydration procedure, another fixation on ice was performed. After washing in PBS, samples were postfixed in 1% OsO_4_ in double distilled water, washed again and dehydrated in a graded series of ethanol (50, 75, 90, 98, and three times 100%) on ice followed by critical point drying out of dry ethanol (CPD 931, Tousimis, USA). The resulting samples were mounted on SEM aluminium stubs, fixed on conductive carbon tape and then sputter-coated with 4nm of platinum/palladium (CCU-10, Safematic, CH). Acquisition of images was performed using In-lens and Everhart-Thornley secondary electron (SE) signals at a working distance of 4mm with a scanning electron microscope (Merlin FE-SEM, Zeiss, DE), operated at an accelerating voltage of 1.5kV.

## Results

### Identification of a Brown Swiss bull with poor semen quality

We examined historic semen quality records from a semen collection centre in Switzerland as part of our ongoing efforts to investigate inherited variation in male reproduction in Brown Swiss bulls [1],[4],[32]. Among 1343 Brown Swiss bulls that produced 70,990 ejaculates, we identified seven bulls from which all ejaculates (between 5 and 28 per bull) were rejected due to immotile (asthenozoospermia), morphologically abnormal (teratozoospermia), or absent (azoospermia) sperm, or a combination thereof. We subsequently focus on one of these bulls (born in 2003) from which preserved hair roots provided a source of DNA for genetic investigations.

Twelve ejaculates from this bull were collected between 15 and 18 months of age. The ejaculates had higher volumes (6.6±1.5 *vs*. 4.7±2.0 ml) and lower sperm concentration (0.2±0.2 *vs*. 1.3±0.5 × 10^9^ sperm cells per ml) than observed in other Brown Swiss bulls of the same age. Almost all (99%) sperm were immotile and had head and tail abnormalities. The proportion of viable sperm was not recorded. Such findings are commonly referred to as oligoasthenoteratozoospermia.

### Candidate causal variants

The genome of the bull that produced ejaculates of insufficient quality (sequencing read archive accession: SAMEA6272098) was sequenced to 11.8-fold coverage using paired-end libraries. Sequence variant genotypes from SAMEA6272098 were compared to those of 397 fertile artificial insemination bulls from various breeds that had average sequencing coverage of 10.0±5.7 fold (between 3.1- and 56.8-fold).

We hypothesized that the drastically reduced semen quality of SAMEA6272098 was due to a recessively inherited deleterious allele. From a catalogue of 41,659,308 autosomal sequence variants, we retained 1655 that were homozygous in SAMEA6272098 but not seen in the homozygous state in the fertile control bulls (**Additional file 4 File S4**). Only three compatible variants were predicted to be deleterious to protein function: a nonsense variant in *MLNR* encoding motilin receptor, a frameshift variant in *QRICH2* encoding glutamine rich 2, and a nonsense variant in *ENSBTAG00000023270* encoding an uncharacterized protein (**Table 1**). Another nine (Ensembl) and ten (Refseq) compatible variants were predicted to have moderate impacts on the respective proteins.

**Table 1:** High impact variants compatible with recessive inheritance.

| Chr | Pos | Ref | Alt | Gene | Transcript abundance in testis tissue (TPM) | Impact on protein | ensembl / refseq / both |
| --- | --- | --- | --- | --- | --- | --- | --- |
| 12 | 18859636 | C | A | *MLNR* | 0.07 ± 0.04 | Cys587Ter | Both |
| 19 | 55436705 | TC | T | *QRICH2* | 31.43 ± 7.35 | Cys1644AlafsTer52 | Both |
| 24 | 10181719 | G | A | *ENSBTAG00000023270* | 0.04 ± 0.03 | Trp248Ter | Ensembl |

In testis transcriptomes from 76 mature bulls, *MLNR* and *ENSBTAG00000023270* were expressed with less than 0.1 transcripts per million (TPM). However, *QRICH2* was transcribed in high abundance (31.43 TPM). Moreover, *QRICH2* shows an extremely testis-biased expression in mammals (e.g., https://gtexportal.org/home/, http://cattlegeneatlas.roslin.ed.ac.uk/) [33]. Pathogenic alleles in human and mouse *QRICH2* cause sperm with multiple morphological abnormalities of the flagella [34],[35],[36]. Thus, we considered *QRICH2* as a compelling functional candidate gene for the sperm defect of SAMEA6272098.

### A 1-bp deletion in the coding sequence of *QRICH2* is associated with aberrant semen quality

The candidate causal variant underpinning the sperm defect is a 1-bp deletion in the sixteenth exon of *QRICH2* (BTA19:55436705TC>T, ENSBTAT00000018337.1:c.4929del). The deletion of a cytosine residue is predicted to alter the reading frame from amino acid 1644 onwards eventually inducing a premature termination codon (ENSBTAP00000018337.1:p.Cys1644AlafsTer52). This leads to a truncated protein that lacks 131 amino acids (7%) from the C-terminal region unless the transcript is subjected to nonsense-mediated mRNA decay.

The 1-bp deletion resides within an approximately 3.625 Mb segment (between 55 and 58.625 Mb) of very low heterozygosity (**Figure 1**). This pattern suggests homozygosity for a haplotype encompassing the 1-bp deletion that SAMEA6272098 inherited from an ancestor that is present in both its maternal and paternal ancestry. The pedigree-derived coefficient of inbreeding of SAMEA6272098 is 7.61%.

**Figure 1:**
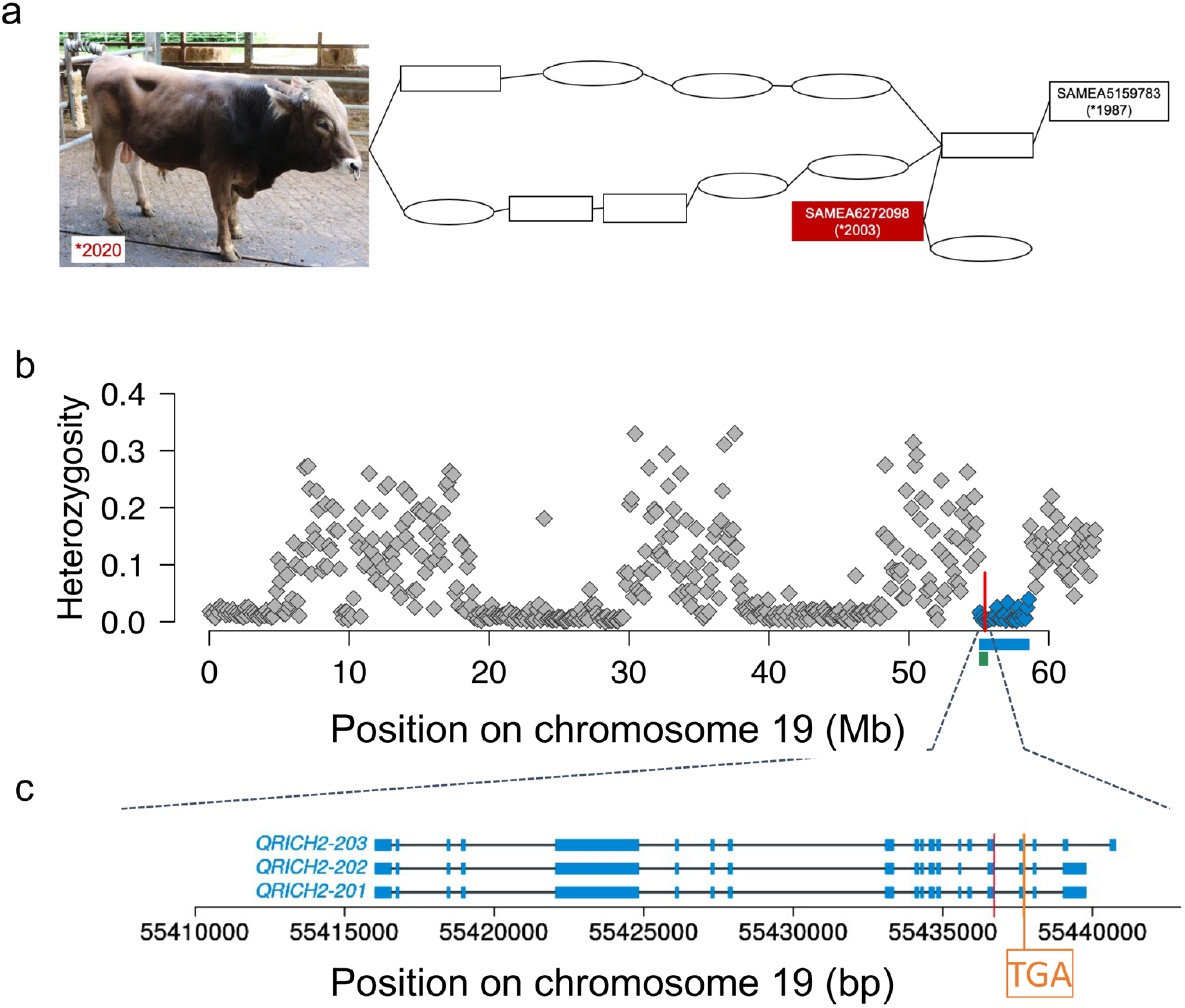
A 1-bp deletion is a candidate causal variant for a sperm morphology defect in Brown Swiss bulls. a) Pedigree of two bulls with a defect of sperm heads and tails. The pedigree only contains suspected carriers of the mutation. Ovals and rectangles represent cows and bulls, respectively. The bull born in 2020 and SAMEA6272098 are related through a common ancestor. SAMEA5159783 is the oldest sequenced mutation carrier in the pedigree. b) Each symbol represents the proportion of heterozygous genotypes observed within a 125 kb window for SAMEA6272098. Blue colour represents a 3.625 Mb segment of extended homozygosity encompassing BTA19:55436705TC>T (red vertical line). The green rectangle indicates the position of a BovineHD-based haplotype that encompasses the 1-bp deletion. c) Structure of bovine *QRICH2* isoforms with transcript-IDs ENSBTAT00000065208, ENSBTAT00000064147, and ENSBTAT00000018337 encoding proteins with 1968 (ENSBTAP00000055962), 1934 (ENSBTAP00000054965) and 1827 (ENSBTAP00000018337) amino acids. Blue rectangles represent exons. The red vertical line indicates the position of the 1-bp deletion (BTA19:55436705TC>T). Orange colour indicates the position of a premature termination codon “TGA” that is introduced due to the shift in translation caused by the 1-bp deletion.

Among 244 and 124 sequenced Brown Swiss and Original Braunvieh cattle, respectively, we detected the 1-bp deletion in the heterozygous state in 25 and 3 individuals, corresponding to an allele frequency of 5.1% and 1.2%. The heterozygous Original Braunvieh bulls were from Germany for which we cannot rule out that the variant was recently introgressed through cross-breeding with Brown Swiss ancestors. The mutation did not segregate among 110 sequenced Original Braunvieh cattle from the Swiss herdbook population. We did not detect the mutation in breeds other than Brown Swiss and Original Braunvieh.

In a catalogue of variants that was established by the 1000 Bull Genomes Project consortium for 3934 cattle from various breeds, the 1-bp deletion was discovered in 16 animals from the Brown Swiss and one animal from the Nordic Red Dairy Cattle breeds. The carrier animal from the Nordic Red Dairy Cattle breed had an estimated proportion of 34% Brown Swiss ancestry. In the 1000 Bull Genomes dataset, only SAMEA6272098 carried the deletion in the homozygous state.

*QRICH2* mRNA was less abundant in six heterozygous carriers of the frameshift variant than 70 bulls that were homozygous for the wild-type allele (23.29 ± 6.76 vs. 32.13 ± 7.01 TPM). Differential expression between heterozygous and wild-type bulls was evident for all *QRICH2* exons (**Additional file 5 File S5**). We observed allelic imbalance at the 1-bp deletion in the total RNA sequencing alignments of all six heterozygous bulls; an average of 292 ± 143 sequencing reads overlapped with the position of BTA19:55436705TC>T, but less than a third (81 ± 32) supported the deletion (**Additional file 5 File S5**), indicating that the RNA is depleted for the transcript carrying the deletion.

### Identification of homozygous haplotype carriers in the Brown Swiss population

To validate the suspected sperm defect arising from the 1-bp deletion, we sought to examine ejaculates from homozygous bulls. Because SAMEA6272098 had been slaughtered in 2005 and semen samples had not been preserved, it was not possible to re-examine its sperm defect. In order to identify homozygous mutation carriers, we employed a dataset that contained 25,243 Brown Swiss and 5,228 Original Braunvieh cattle with phased Illumina BovineHD Bead chip genotypes at 17,579 SNPs located on bovine chromosome 19. These animals were genotyped for routine genomic breeding value prediction in Switzerland.

Of the 30,471 genotyped Brown Swiss and Original Braunvieh cattle, 285 (28 females, 257 males) also had whole-genome sequence-called genotypes at the BTA19:55436705TC>T variant. Twenty-one sequenced animals (1 female, 20 males) were heterozygous carriers of the 1-bp deletion. Under the assumption that these 21 animals inherited the 1-bp deletion from a common ancestor, we screened their phased array-derived genotypes for a shared haplotype encompassing BTA19:55436705TC>T. This approach identified a 675 kb haplotype encompassing 181 Illumina BovineHD BeadChip SNPs and spanning from 54,992,461 to 55,667,539 bp on BTA19 that all 21 heterozygous animals carried in the heterozygous state (**Figure 1, Additional file 2 File S2**).

Of the 264 sequenced animals that did not carry the 1-bp deletion, only one animal (SAMEA7573647) carried the haplotype in the heterozygous state. This animal was a fertile paternal half-sib of SAMEA6272098. Upon manual inspection of the read alignments, we noticed that only one properly mapped sequencing read overlapped with BTA19:55436705TC>T in that bull (**Additional file 6 File S6**). This read supported the 1-bp deletion, despite the bull being genotyped as homozygous for the reference cytosine residue. Low sequencing coverage likely resulted in the undercalling of heterozygous genotypes. Thus, the microarray-derived haplotype was in full concordance with the genotypes at the BTA19:55436705 TC>T variant in the 285 sequenced animals.

The 675 kb haplotype had a frequency of 5.4% among 25,243 genotyped Brown Swiss cattle. Haplotype frequency didn’t change notably over the past 30 years. We detected 2712 heterozygous and 54 and homozygous haplotype carriers. Since 79 homozygous animals were expected assuming random mating, the observed haplotype distribution deviates slightly from Hardy-Weinberg proportions (P=0.03). The maternal grandsire from the heterozygous bull from the Nordic Red Dairy Cattle breed carried the haplotype in the heterozygous state. Of the homozygous haplotype carriers, 35 were females and 19 were males. None of the homozygous bulls had been selected as an artificial insemination sire. Thus, their semen quality was not examined. Of the 35 homozygous females, 33 gave birth to at least one offspring, indicating that their fertility was not compromised. Inspection of uncorrected milk records in heterozygous cows indicated that their milk production is normal. None of the 5,228 Original Braunvieh cattle carried the 675 kb haplotype.

We used semen quality data from 1133 Brown Swiss bulls that were previously analyzed by Hiltpold et al. ([1],[4]) to investigate if the haplotype is associated with semen quality in the heterozygous state. We examined an average number of 55 (between 5 and 242) and 58 (between 2 and 573) ejaculates, respectively, from 143 heterozygous haplotype carriers and 990 bulls that did not carry the haplotype. Average ejaculate volume (in ml), sperm concentration (million sperm per ml), and percentage of motile sperm was 4.68, 1263, and 85 in heterozygous haplotype carriers and 5.02, 1218, and 84 in non-carriers, respectively. The proportion of ejaculates that were rejected or contained an excess of sperm with head and tail anomalies did not differ notably between heterozygous carriers and non-carriers, supporting a recessive mode of inheritance.

All male homozygous haplotype carriers had been slaughtered before our study was conducted, preventing ejaculate collection and semen analyses. In order to identify homozygous bulls for phenotypic investigations, we provided the coordinates of the 675 kb haplotype to the Brown Swiss genetic evaluation centres in Switzerland, Austria and Germany. A haplotype screen in the genomic selection reference populations identified a six-month-old bull born in 2020 that carried the haplotype in the homozygous state. The sire of SAMEA6272098 was in the maternal (6^th^ ancestral generation) and paternal (5^th^ ancestral generation) ancestry of the six-month-old homozygous haplotype carrier indicating inbreeding due to an obligate carrier of the 1-bp deletion (**Figure 1a**). Sanger sequencing confirmed homozygosity for the 1-bp deletion (BTA19:55436705TC>T). The bull appeared regularly developed and healthy.

### Ejaculates of a homozygous bull contain immotile sperm with multiple morphological abnormalities

We kept this bull at a research station together with other bulls of similar age. At 14 months, scrotal circumference was relatively small (31 cm) possibly indicating delayed puberty. The first ejaculate collected from the bull had a volume of 5 ml, but it contained only 0.03 × 10^9^ sperm per ml (**Table 2**). None of the sperm were motile and 97.3% had tail or head abnormalities. Scrotal circumference increased with age (34.5 cm at 20 months) but remained relatively small compared to other bulls. We collected six additional ejaculates from the bull at 16, 17, and 20 months. The ejaculates had an average volume of 4.0±0.9 ml and contained 0.2±0.1 × 10^9^ sperm per ml, confirming normal volume but aberrantly low sperm concentration and sperm count. More than 99% of the sperm had major defects and less than 1% were motile. Histological examination of the testis after slaughter indicated that Leydig cells, rete testis, and tubuli seminiferi were formed normally and spermatogenesis was detectable.

**Table 2:**
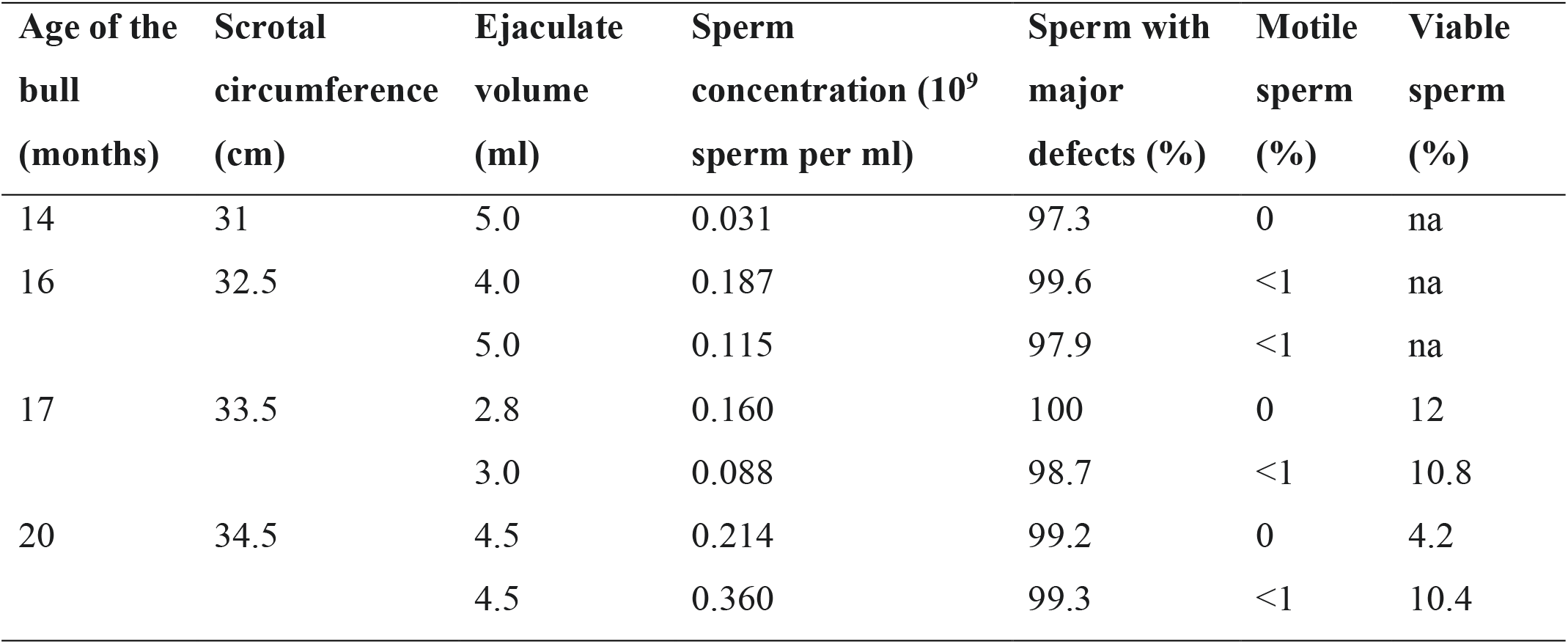
Semen analysis in a bull homozygous for the 1-bp deletion in the coding sequence of *QRICH2*.

| Age of the bull (months) | Scrotal circumference (cm) | Ejaculate volume (ml) | Sperm concentration (10<sup>9</sup> sperm per ml) | Sperm with major defects (%) | Motile sperm (%) | Viable sperm (%) |
| --- | --- | --- | --- | --- | --- | --- |
| 14 | 31 | 5.0 | 0.031 | 97.3 | 0 | na |
| 16 | 32.5 | 4.0 | 0.187 | 99.6 | <1 | na |
|  |  | 5.0 | 0.115 | 97.9 | <1 | na |
| 17 | 33.5 | 2.8 | 0.160 | 100 | 0 | 12 |
|  |  | 3.0 | 0.088 | 98.7 | <1 | 10.8 |
| 20 | 34.5 | 4.5 | 0.214 | 99.2 | 0 | 4.2 |
|  |  | 4.5 | 0.360 | 99.3 | <1 | 10.4 |

Microscopic semen analysis revealed multiple morphological abnormalities in almost all sperm primarily affecting the flagella (**Figure 2, Additional file 7 File S7**). Morphological abnormalities included shortened, coiled, bent, looped, thickened, doubled and absent tails. Loose tails were not detected. Approximately 20% of the examined sperm were either underdeveloped, had irregularly shaped heads or nuclear vacuoles. Eosin-nigrosin staining indicated that between 4 and 12% of the sperm were viable despite morphological abnormalities of the head and flagella (**Figure 2c**).

**Figure 2:**
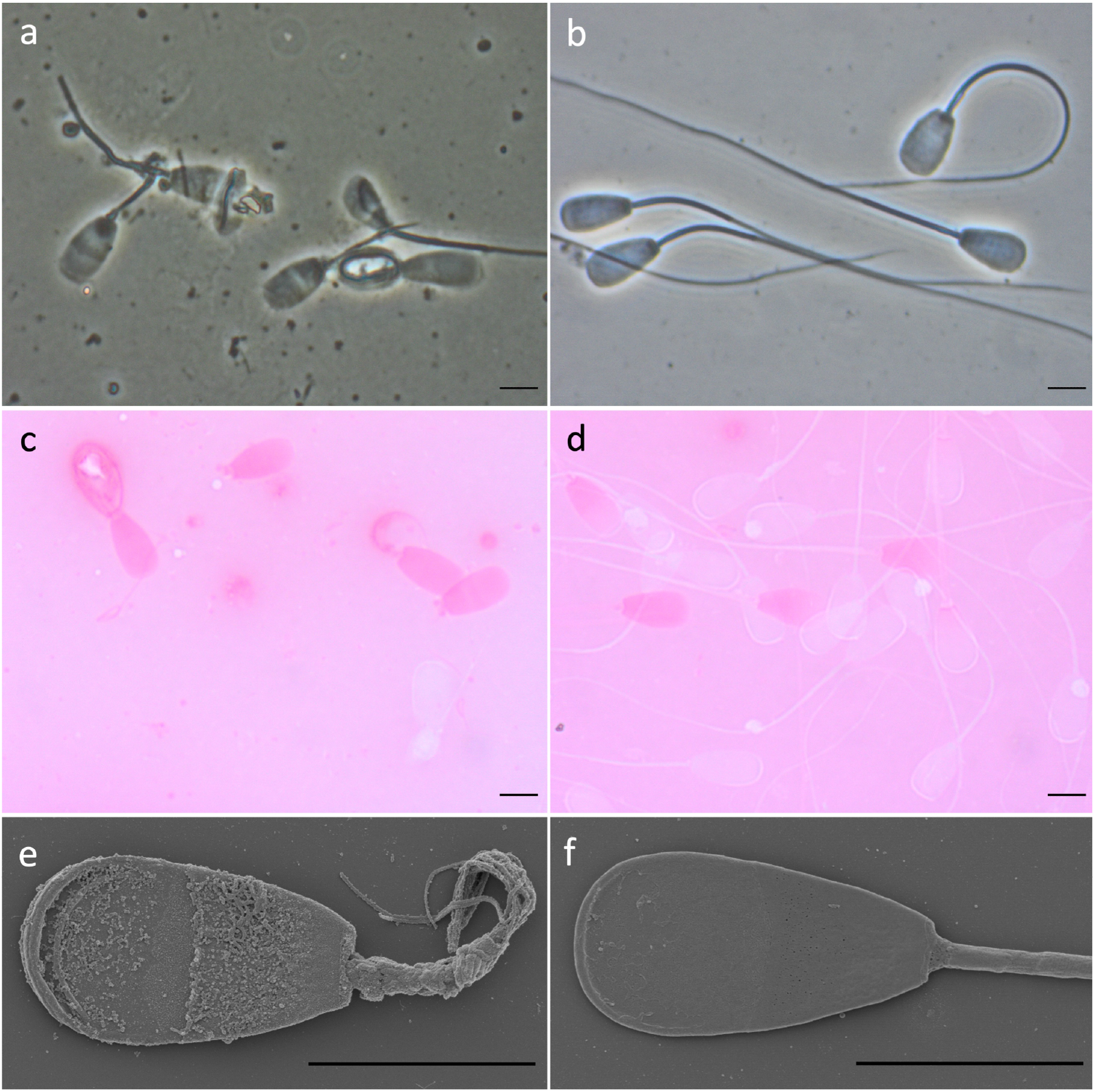
Sperm of a bull homozygous for the 1-bp deletion and of a control bull. Representative phase-contrast (a, b), eosin-nigrosin stained (c, d), and scanning electron microscopy images (e, f) of sperm of a bull homozygous for the 1-bp deletion (a, c, e) and of a control bull (b, d, f) with normal sperm quality. Sperm of the affected bull show multiple morphological abnormalities of head and flagella (a). Phase-contrast microscopy also revealed numerous cell debris particles. Viable sperm remain unstained and appear white in eosin-nigrosin stained images whereas dead sperm are stained and appear purple (c, d). Flagella of the affected bull are irregularly shaped and mostly have an uneven surface (e). Scale bar: 5µm

Transmission electron microscopy cross-sections of sperm from the affected bull confirmed multiple ultrastructural defects of the flagellum and head (**Figure 3, Additional file 8 File S8**). The axonemes of most flagella deviated from the typical assembly of nine outer microtubule doublets surrounding the central pair. A variable number of outer microtubule doublets was missing in the principal and end piece region of most examined flagella. We also detected sperm with multiple flagellar structures within the same cell membrane, indicating that the flagellum was folded around the head instead of being elongated which is typical for underdeveloped sperm (**Additional file 8 File S8**). Occasionally, we found axonemes in cross-sections of the principal piece that appeared regularly organized.

**Figure 3:**
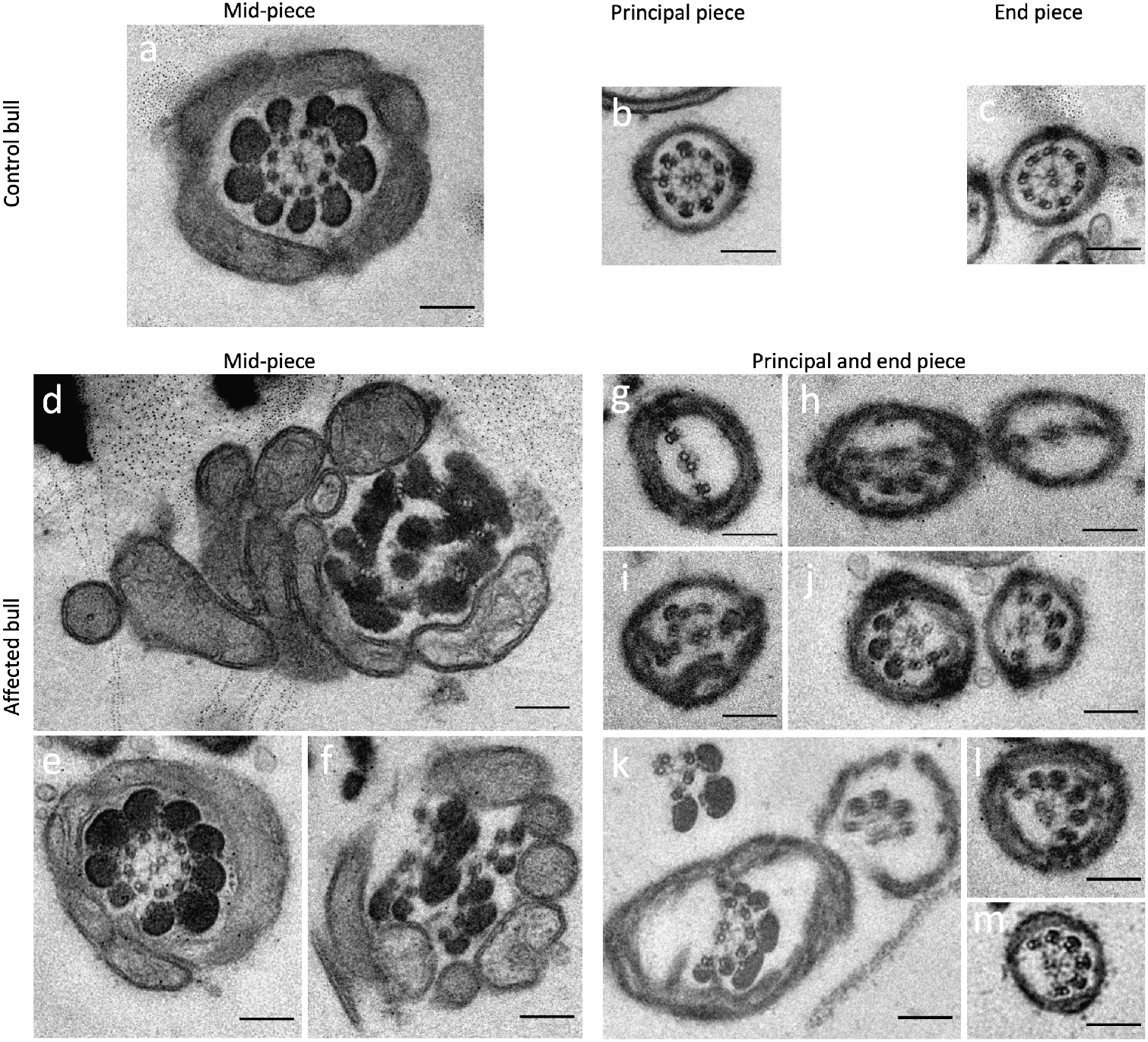
Transmission electron microscopy cross-sections of bovine sperm flagella. Representative TEM cross-sections of sperm flagella at the mid-piece, principal piece and end piece in a control bull (a-c) and a bull homozygous for the 1-bp deletion in *QRICH2* (d-m). Irregularities in the flagellar cross-sections from the homozygous bull prevent unambiguous assignment of the principal and end piece (g-m). The axonemes of most flagella lack some of the outer microtubules resulting in deviations from the typical arrangement of nine outer microtubule doublets surrounding the central pair in the homozygous bull. Cross-sections also show irregular assembly of axonemes and structures such as outer dense fibres that are supposed to enclose the axonemes (d, f). Scale bar: 200nm.

## Discussion

We detect a recessive, frameshift-inducing, 1-bp deletion in *QRICH2* in two Brown Swiss bulls producing ejaculates with low sperm concentration and immotile sperm with multiple morphological abnormalities. Semen samples from the bulls were not used for artificial inseminations because this disorder very likely prevents fertilization *in vivo* due to the inability of the sperm to move. Intracytoplasmic sperm injection (ICSI) – albeit not a common practice in cattle breeding – could possibly enable fertilization because some sperm were viable and had normally shaped heads. In fact, ICSI enabled fertilization using sperm with multiple morphological abnormalities of the flagella due to deleterious variants in *QRICH2* and normal offspring development in humans [36].

We applied an indirect haplotype test to estimate that the 1-bp deletion segregates at a frequency of 5% in the Brown Swiss population. Assuming random mating, the deletion is expected to occur in the homozygous state in 1 out of 400 bulls. However, the proportion of homozygous bulls was lower in our study. None of 1133 Brown Swiss bulls for which we had microarray-derived genotypes and routinely assessed semen quality phenotypes, carried the haplotype in the homozygous state. The 1-bp deletion resides on a haplotype that is inherited from a common ancestor. The mating of closely related individuals is generally avoided to prevent inbreeding, and therefore haplotypes may occur less frequent in the homozygous state than expected under random mating. In principle, the frequency of the 1-bp deletion may be lower than estimated from an indirect haplotype test. If a mutation occurred relatively recently, the haplotype encompassing the mutation may be indistinguishable from its ancestral form that does not carry the mutation. In such a situation, a haplotype-based test overestimates the frequency of the mutation [37]. However, we didn’t find evidence that an ancestral version of the haplotype segregates in the Brown Swiss population, as the 675 kb haplotype was in full concordance with the 1-bp deletion. Moreover, the haplotype frequency agreed well with the frequency of the 1-bp deletion estimated from whole-genome sequence variants.

The birth years of the two homozygous bulls span almost two decades. Only one of them was selected as a potential artificial insemination sire. Since immotile sperm with multiple morphological abnormalities became apparent during routine semen analyses, all ejaculates from this bull were discarded and not used for artificial inseminations. Heterozygous bulls produce normal sperm. If the semen of prospective artificial insemination and natural mating bulls is examined, this sperm defect can be recognized easily and won’t manifest in repeated breedings and low non-return rates as it is the case for infertility disorders that are not apparent from standard semen analyses [8]. Economic losses due to this defect are negligible in the Brown Swiss cattle population because the frequency of the 1-bp deletion is low and aberrant semen quality of homozygous bulls can be detected easily.

The widespread use of individual mutation carriers in artificial insemination can result in the frequent manifestation of recessive alleles within short time [38]. It is advised to monitor the BTA19:55436705TC>T variant in the Brown Swiss population, e.g., using customized microarrays, and implement genome-based mating programs to avoid the frequent emergence of bulls that produce ejaculates containing immotile sperm. We did not detect the BTA19:55436705TC>T variant in Swiss Original Braunvieh animals suggesting that the mutation occurred relatively recently possibly after the bifurcation of the Braunvieh breed into a dairy (Brown Swiss) and dual-purpose (Original Braunvieh) lineage. We found the 1-bp deletion in one animal from the Nordic Red Dairy Cattle breed likely resulting from crossbreeding with a heterozygous Brown Swiss sire [39]. Thus, our study adds to an increasing number of alleles with undesired effects that had been introgressed from foreign breeds into the Nordic Red Dairy Cattle breed [40],[41],[42].

Human and mouse orthologs of *QRICH2* harbour loss-of-function variants that result in multiple morphological abnormalities and ultrastructural defects of the sperm flagella, thereby leading to immotile sperm and infertility *in vivo* [34],[35],[43]. We replicate these findings in ejaculates from two bulls homozygous for a frameshift-inducing 1-bp deletion in bovine *QRICH2* suggesting evolutionarily conserved protein function. The 1-bp deletion is predicted to induce a premature termination codon that shortens the protein by 131 amino acids. The truncated protein may be retained with reduced functionality or the transcript may be degraded via nonsense-mediated mRNA decay. The position of the premature termination codon complies with both canonical and non-canonical rules for nonsense-mediated mRNA decay [44],[45],[46]. Allelic imbalance at the 1-bp deletion and lower levels of *QRICH2* mRNA in testis transcriptomes of heterozygous than wild-type bulls confirms that the premature termination codon triggers nonsense-mediated mRNA decay. These findings show that the 1-bp deletion constitutes a loss-of-function allele of bovine *QRICH2*, thereby supporting the pathogenicity of a predicted deleterious variant residing in a similar domain of human *QRICH2* [35].

So far, QRICH2 has primarily been implied in sperm flagellar formation. Apart from flagellar abnormalities, we noticed sperm head anomalies, drastically reduced sperm concentration and low sperm count in all ejaculates examined from bulls homozygous for the 1-bp deletion. Sperm concentration and total sperm count were also at the lower bound of the normal range in humans with deleterious *QRICH2* alleles [34],[35],[43]. These findings suggest that loss of QRICH2 functionality not only compromises the assembly of sperm flagella, but generally impairs spermiogenesis. Variants other than the 1-bp deletion may contribute to the aberrant semen quality of the two examined Brown Swiss bulls. However, pinpointing putative modifier loci is difficult for rare alleles [47] and was not attempted for the 1-bp deletion in *QRICH2*.

To the best of our knowledge, this is the first time that consequences arising from a loss-of-function allele in *QRICH2* are described in a species other than mice and human. Apart from producing anomalous ejaculates, homozygous bulls are indistinguishable from wild-type and heterozygous individuals. Homozygous cows were not examined clinically in our study, but their undisturbed fertility and normal milk production likely indicate an overall normal health [48]. Moreover, no QTL for economically relevant traits have been detected next to BTA19:55436705TC>T in Brown Swiss cattle [49]. These findings suggest that the 1-bp deletion does not compromise traits other than semen quality. This agrees with loss-of-function alleles in other genes with extreme testis-biased expression [8],[9],[10],[50], and corroborates findings in humans and mice with loss-of-function alleles in *QRICH2* [34] suggesting that QRICH2 is dispensable for somatic development in mammals. The deviation of the haplotype encompassing the 1-bp deletion from Hardy-Weinberg proportions (P=0.03) is likely due to either selective breeding or the inability of homozygous males to contribute to the next generation rather than fatal consequences arising from homozygosity.

Our analyses in two homozygous bulls provide evidence for the causality of the 1-bp deletion from a statistical, functional, and comparative genomics point of view [51],[52]. We discovered the deletion using a phenotype-driven approach in a historic sample and verified its phenotypic consequences in a second bull born almost two decades later that we found through a genotype-driven screen. Transcriptome analyses show that the 1-bp deletion triggers nonsense-mediated mRNA decay, corroborating it constitutes a loss-of-function allele. Orthologs of bovine *QRICH2* harbour loss-of-function alleles, some of them residing in the same domain as the 1-bp deletion identified in our study, that cause sperm defects mirroring those of the homozygous Brown Swiss bulls, suggesting QRICH2 is essential for male fertility in mammals. While a formal proof for the causality remains to be produced, these pieces of evidence are sufficient to warrant monitoring of the 1-bp deletion in cattle breeding programs.

## Conclusions

A 1-bp deletion in the coding sequence of bovine *QRICH2* is likely causal for low sperm concentration and immotile sperm with multiple morphological abnormalities in the homozygous state. The 1-bp deletion has a frequency of 5% in the Brown Swiss cattle population. Homozygous bulls are unsuitable for breeding as they are very likely infertile *in vivo*. Apart from poor semen quality, homozygous bulls are indistinguishable from heterozygous and wild-type animals. Immediate economic consequences arising from the undesired allele are negligible. The monitoring of the defective allele in the Brown Swiss cattle population using customized genotyping is advised to avoid an increase in allele frequency.

## Declarations

### Ethics approval and consent to participate

Our study was approved by the veterinary office of the Canton of Zurich, Switzerland (animal experimentation permit ZH 181/19).

### Consent for publication

Not applicable

### Availability of data

Whole-genome sequencing data of 397 fertile bulls and the infertile bull are available at the European Nucleotide Archive (ENA) of the EMBL under sample accessions listed in **Additional file 1 File S1**. Whole-genome sequencing data of 285 cattle used to identify a trait-associated haplotype are available at the European Nucleotide Archive (ENA) of the EMBL under sample accessions listed in **Additional file 1 File S1**. DNA and RNA sequencing data of 76 bulls of the eQTL cohort are available at the European Nucleotide Archive (ENA) of the EMBL under sample accessions listed in **Additional file 3 File S3**.

## Author’s contributions

HP conceived the study. MH and HP analysed the data. MH and FJ analysed semen samples of the homozygous bull. NKK called sequence variant genotypes. XMM established the eQTL cohort. ZHF carried out sequencing experiments. HP, HS and FRS analysed genotyping data from the Brown Swiss reference populations. MS organised genetic material for sequencing. UW provided semen quality data. HP and MH wrote the manuscript. All authors read and approved the final manuscript.

## Acknowledgements

We thank Dr. Franz Birkenmaier (Allgäuer Herdbuchgesellschaft) for support in identifying homozygous bulls as well as Dr. Melissa Terranova and Flavio Ferrari (AgroVet-Strickhof) for the handling of the bull. We thank Braunvieh Schweiz for providing genotype data. We are thankful for the excellent technical support provided by the ETH Zürich technology platform FGCZ (https://fgcz.ch/) concerning DNA and RNA sequencing. We are grateful to Dr. Cecilia Bebeacua and Stephan Handschin from the ETH Zürich technology platform ScopeM (https://scopem.ethz.ch/) for outstanding support in electron microscopy. We appreciate support from veterinarians from the meat inspection and the slaughterhouse staff in sampling of bull testes as well as Till Graf, Patrick Fitzi, and Remo Hengartner for their contribution to collect tissue.

## Additional Files

### Additional file 1 File S1

File format: xlsx

Title: Sequence read archive accessions for the sequenced animals.

Description: The spreadsheet file contains accessions from the European Nucleotide Archive that point to sequencing data for cattle from various breeds. The data in the file are organised in three spreadsheets. The accession for the bull that produced ejaculates with poor quality is listed in «case_SRA_ID». Accessions for 397 fertile control animals from various breeds are listed in «controls_SRA_IDs». Accessions, sequence-based genotypes and haplotypes for 285 cattle used to assign the 1-bp deletion onto a haplotype are listed in «array-based genotypes».

### Additional file 2 File S2

File format: txt

Title: Coordinates of a diagnostic haplotype.

Description: Illumina BovineHD coordinates (SNP-name, chromosome, physical position [ARS-UCD1.2]) and haplotype allele of 181 markers that define the diagnostic 675 kb haplotype.

### Additional file 3 File S3

File format: xlsx

Title: Accessions for the eQTL cohort.

Description: Accessions for sequencing reads obtained from RNA and DNA prepared from testis tissue of 76 mature bulls.

### Additional file 4 File S4

File format: xlsx

Title: Candidate causal variants.

Description: Functional consequences predicted for 1655 variants compatible with recessive inheritance based on the Ensembl (version 104) and Refseq (version 106) annotation of the bovine genome.

### Additional file 5 File S5

File format: png

Title: *QRICH2* mRNA analysis.

Description: a) Exon-specific expression (quantified in transcripts per million [TPM]) for *QRICH2* in testis tissue of six heterozygous (green) and 70 homozygous wild-type (grey) bulls. b) Integrative Genomics Viewer coverage tracks from RNA sequence read alignments overlapping the frameshift-inducing 1-bp deletion (red arrow) in six heterozygous bulls. The identifiers of the coverage tracks refer to accessions from the sequence read archive.

### Additional file 6 File S6

File format: png

Title: DNA sequence alignment of SAMEA6272098.

Description: Output from «samtools tview» centered on BTA19:55436705TC>T representing DNA sequence read alignments from a bull (SAMEA6272098) that carries the 675 kb haplotype in the heterozygous state, but was genotyped as homozygous for the reference allele at position of the BTA19:55436705TC>T variant. The asterisk within the only read overlapping BTA19:55436706 indicates that the bull carries the 1-bp deletion.

### Additional file 7 File S7

File format: tif

Title: Phase-contrast images of sperm from a bull homozygous for the 1-bp deletion. Description: All images display sperm with major sperm head and/or flagellar defects. Very short, shortened or absent flagella were the most prevalent flagellar abnormalities: very short flagella (a, b, d, e, f, g, i, k, s, u), shortened flagella (c, e, h, i, l, q, r,), and absent flagella (c, i, j, o, t). Short doubled (e, k, s) or short thickened (e, k, i, n, q) were apparent too. Some flagella were strongly folded (m, y right sperm) or coiled (y left sperm). Many sperm showed defective heads additionally to the defective flagella: pyriform (c, i, m, s, t), round (g, r), abnormal contour (p), and diadem defect/vacuoles (b, o, q, u, y). Underdeveloped sperm with the flagella strongly folded around the sperm head are considered the most severe sperm defects in bull (d right sperm, t right sperm, v, w, x). Scale bar: 5µm.

### Additional file 8 File S8

File format: tif

Title: TEM and SEM images of sperm from a bull homozygous for the 1-bp deletion. Description: TEM cross-section of an underdeveloped sperm with multiple flagellar structures next to the nucleus (black) enclosed by a cell membrane (a). Underdeveloped sperm with flagellum curled on head visualized using SEM (b). Longitudinal TEM cross-section (c) and SEM (d) of sperms with a thickened vesicularized structure at the sperm neck and multiple disorganized flagellar structures at the mid-piece. Longitudinal TEM cross-section of the neck region of a normal sperm from a control bull (e). Scale bar: 1µm.

