## Supplementary figures and images for "A 1-bp deletion in bovine *QRICH2* causes low sperm count and immotile sperm with multiple morphological abnormalities"

### Supporting File 5

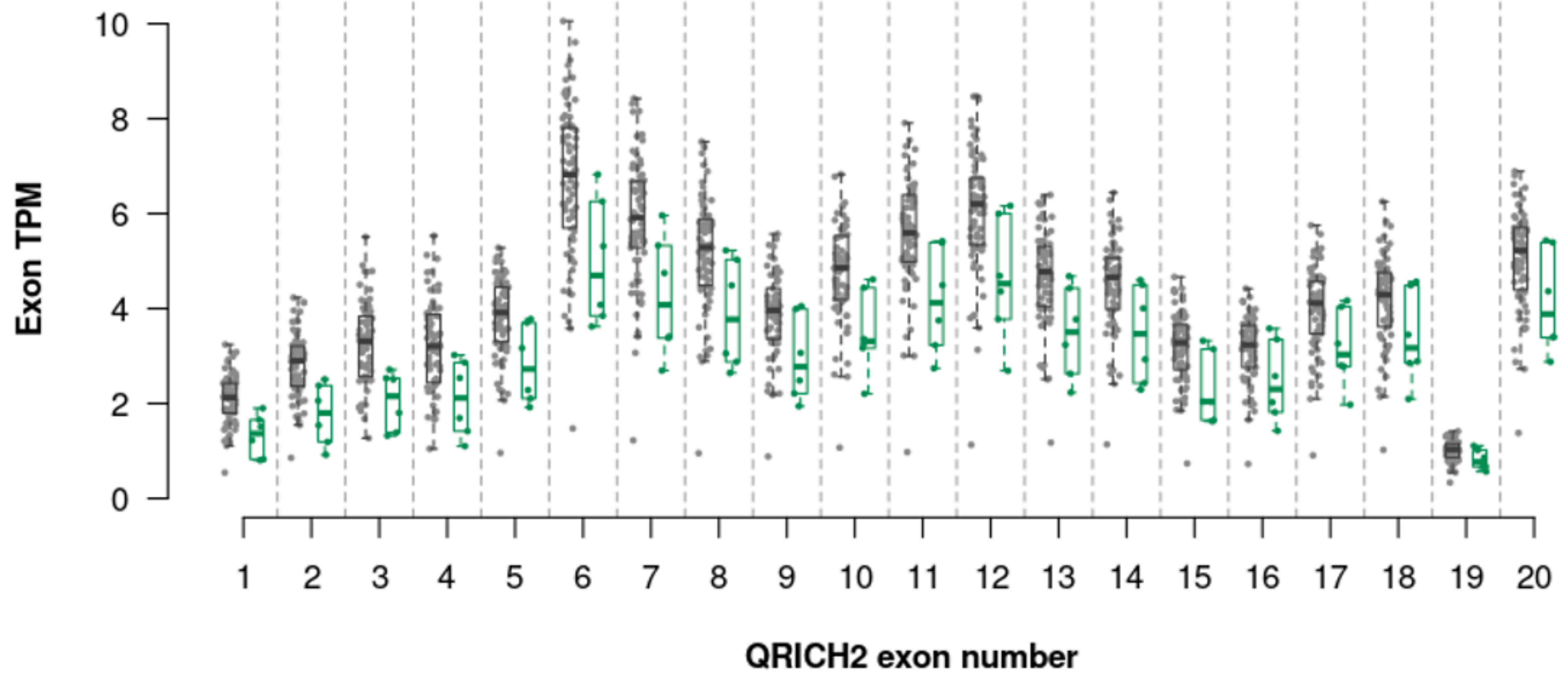

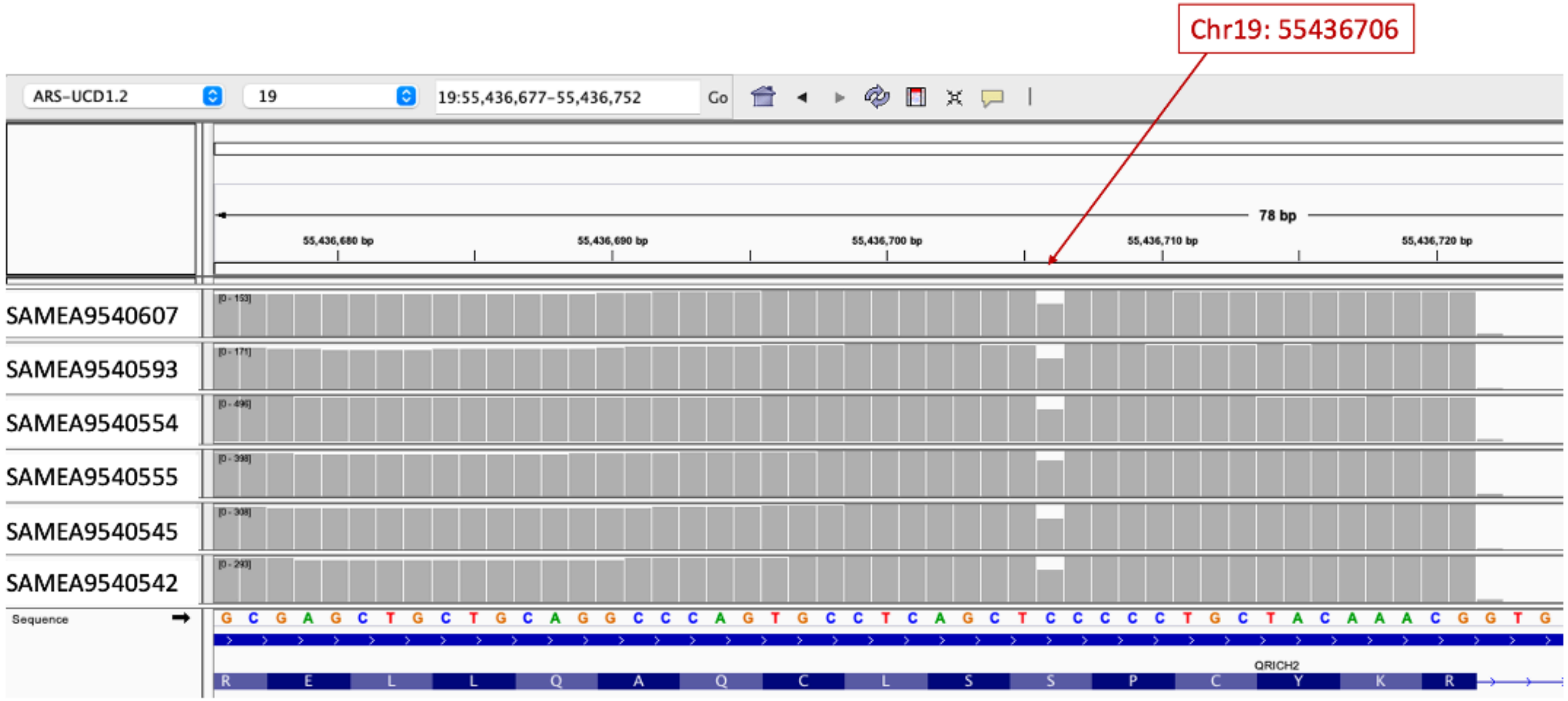

### Supporting File 7

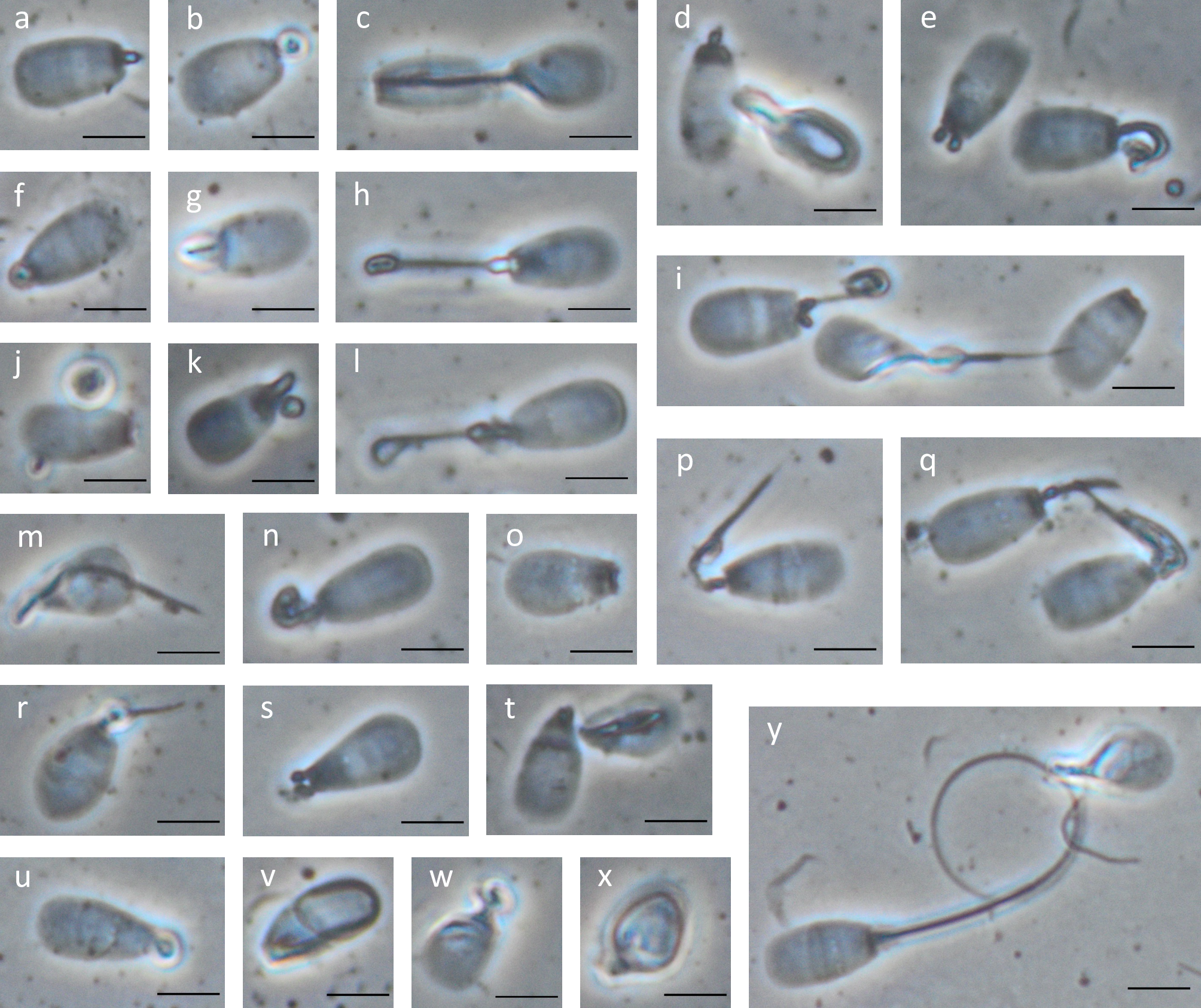

### Supporting File 8

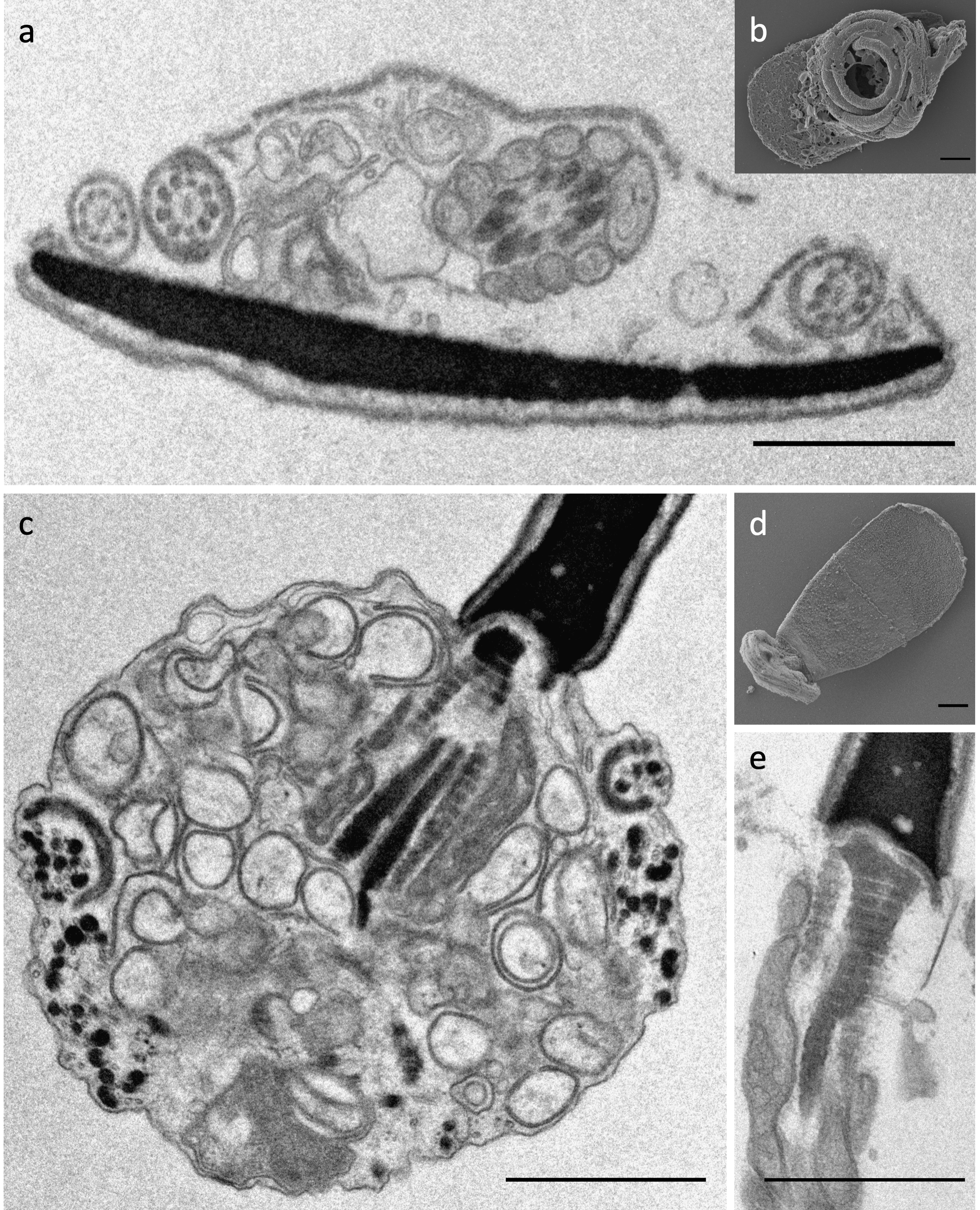
