## Supporting File 6 for "A 1-bp deletion in bovine *QRICH2* causes low sperm count and immotile sperm with multiple morphological abnormalities"

```
661  55436671  55436681  55436691  55436701  55436711  55436721  55436731  55436741  55436751
CAGATGATCCGCGAGCTGCTGCAGGCCCAGTGCCTCAGCTCCCCCTGCTACAAACGGTGGGCAGCCGGGCTGGGCCTGGGAGCCTGGCACCAG
........................................K................                  ..................
........................................*................
```
